# The rice ethylene receptor OsERS1 negatively regulates the shoot growth and salt tolerance in rice seedlings

**DOI:** 10.1101/2025.01.15.633154

**Authors:** Shigeto Morita, Soma Tanaka, Yuzuki Tani, Jun’ichi Nakamura, Masa H. Sato, Shigeru Satoh, Takehiro Masumura

**Author notes:** Correspondence: Shigeto Morita, Laboratory of Genetic Engineering, Graduate School of Life and Environmental Sciences, Kyoto Prefectural University, Shimogamo, Sakyo, Kyoto 606-8522, Japan.

## Abstract

Ethylene is a gaseous plant hormone that regulates various aspects of growth and development. It promotes shoot growth in etiolated rice seedlings, which is crucial for seedling establishment and has significant agricultural implications. Of the five ethylene receptors in rice, *OsERS1* is the most abundantly expressed in seedlings. Therefore, we investigated the localization and role of OsERS1 in shoot growth. When expressed in rice protoplasts, OsERS1 was localized in the endoplasmic reticulum. Knockout mutants of *OsERS1* exhibited enhanced shoot growth in etiolated seedlings under normal and ethylene-treated conditions. Additionally, the induction of ethylene-responsive genes was increased in the mutants, indicating an elevated ethylene response due to *OsERS1* knockout. The mutants also showed increased salt tolerance. In conclusion, our findings suggest that OsERS1 negatively regulates shoot growth and salt tolerance in etiolated rice seedlings.

## Introduction

Ethylene is a gaseous plant hormone that regulates various aspects of growth and development, such as seed germination, elongation, fruit ripening, and senescence (Bleecker and Kende 2000). Molecular genetic studies in *Arabidopsis* have elucidated the mechanisms of ethylene perception and signal transduction (Bakshi *et al*. 2015). In *Arabidopsis*, ethylene is perceived by five receptors: ETHYLENE RESPONSE1 (ETR1), ETR2, ETHYLENE INSENSITIVE4 (EIN4), ETHYLENE RESPONSE SENSOR1 (ERS1), and ERS2. These receptors act as negative regulators, suppressing the ethylene response in the absence of the hormone by activating the downstream Raf-like Ser/Thr kinase CONSTITUTIVE TRIPLE RESPONSE 1 (CTR1), which leads to the phosphorylation and degradation of EIN2, a key regulator of the ethylene response. In the presence of ethylene, the receptors and CTR1 are inactivated, resulting in EIN2 dephosphorylation and stabilization. This activates the downstream transcription factors EIN3 and EIN3-LIKE1 (EIL1), triggering the transcription of ethylene-responsive genes.

Ethylene receptors are membrane-bound proteins primarily localized in the endoplasmic reticulum (ER). They share a common structure, including an N-terminal ethylene-binding domain, followed by GAF and His kinase domains. These receptors are classified into two subfamilies: subfamily 1 includes ETR1 and ERS1, while subfamily 2 comprises ETR2, ERS2, and EIN4. Although their functions partially overlap, leading to no observable phenotypes in single loss-of-function mutants (Hua and Meyerowitz 1998), each receptor has unique roles. For example, ETR1 and EIN4 negatively regulate seed germination under salt stress, while ETR2 has a positive role (Wilson *et al*. 2014).

Rice (*Oryza sativa* L.) is an essential staple crop as well as a model of monocotyledonous plants. In etiolated rice seedlings, ethylene suppresses root growth while promoting shoot (coleoptile) elongation (Ma *et al*. 2013). This ethylene-induced shoot elongation is unique to rice and is not observed in other monocots (Yin *et al*. 2023). Since post-germination growth notably affects seedling establishment, understanding the regulatory mechanisms of shoot elongation has agricultural significance.

Ethylene signaling pathways in rice have conserved and divergent features with those in *Arabidopsis* (Zhao *et al*. 2021). There are several orthologous genes to *Arabidopsis* ethylene signaling components in rice, and a linear signaling pathway involving ethylene receptors, OsCTR2, OsEIN2, and OsEIL1/ OsEIL2 is conserved in rice. Additionally, rice possesses an alternative ethylene signaling pathway mediated by MHZ1/OsHK1, functioning parallel to the OsEIN2 pathway (Zhao 2021). Rice has five ethylene receptors (Yau *et al*. 2004): OsERS1 and OsERS2 (subfamily 1) and OsETR2, OsETR3, and OsETR4 (subfamily 2) (Zhao 2021). Since single mutants of rice ethylene receptors show distinct phenotypic changes, each isoform plays different physiological roles. For instance, mutants of *OsERS 2*, *OsETR3*, and *OsERS2* exhibit enhanced ethylene responses in shoots (Wuriyanghan *et al*. 2009), whereas the *OsERS1* mutant shows an increased root ethylene response (Ma *et al*. 2014), indicating that these receptors negatively regulate ethylene responses in rice. Furthermore, some ethylene receptor isoforms influence growth throughout the life cycle. For example, a knockdown line of *OsETR2* displays early flowering and reduced starch accumulation under field conditions (Wuriyanghan 2009). The *OsERS1 OsERS2* double mutant shows severe defects in shoot growth and seed setting, emphasizing the critical roles of these receptors in rice growth regulation (Zhou *et al*. 2020).

Among the five ethylene receptors in rice, *OsERS1* (RAP-DB Gene ID: Os03g0701700) is most abundantly expressed in seedlings (Yau 2004). It also exhibits high expression in various tissues from the vegetative to the reproductive stages, as indicated by the RiceXPro expression profile database (https://ricexpro.dna.affrc.go.jp/). Given this expression pattern, we were interested in the role of *OsERS1* in growth regulation. While OsERS1 has previously been reported to localize on the plasma membrane (Yu, Yau and Yip 2017), differing from most ethylene receptors, we re-examined its localization in this study using a green fluorescent protein (GFP)-fusion protein. Although the involvement of *OsERS1* in the root ethylene response of etiolated seedlings has been established (Ma 2014), its role in the shoot remains unclear. Therefore, we examined the shoot ethylene response of *OsERS1* knockout (KO) mutants created using the CRISPR/Cas9 system in this study. Additionally, we investigated the role of *OsERS1* in salt tolerance, as ethylene also contributes to the regulation of salt stress response (Riyazuddin *et al*. 2020; Zhang *et al*. 2016).

## Materials and Methods

### Plasmid construction for the expression of the OsERS1–GFP fusion protein

To achieve efficient expression of the GFP fusion protein in rice cells, we replaced the Cauliflower mosaic virus (CaMV) 35S promoter in the CaMV35S-sGFP(S65 T)-NOS3′ vector (Chiu *et al*. 1996; provided by Dr. Yasuo Niwa) with a modified rice ubiquitin promoter (derived from pZH2BikU; Kuroda, Kimizu and Mikami 2010) at the *Hin* d III and *Xba* I sites. The resulting plasmid was named OsUbiP-GFP/pUC19. The entire coding sequence (CDS) of *OsERS1* was amplified by PCR from a full-length cDNA clone (accession no. AK067813, provided by NARO DNA Bank, Japan) using the primers listed in Table S1 and a high-fidelity DNA polymerase, KOD Plus (Toyobo, Osaka, Japan). The amplified fragment was inserted into the *Xba* I and *Nco* I sites of OsUbiP-GFP/pUC19, creating the OsERS1–GFP fusion plasmid (Fig. S1).

Additionally, we expressed a red fluorescent protein (RFP) fusion of *Arabidopsis* Sec22 (AtSec22) to serve as a marker for the ER and Golgi apparatus (Chatre *et al*. 2005; Uemura *et al*. 2004). To construct a plasmid for expressing TagRFP-AtSec22, the GFP CDS in OsUbiP-GFP/pUC19 was replaced with the TagRFP-AtSec 22 CDS fragment from a CaMV 35S-TagRFP-AtSec22 fusion plasmid (Morita *et al*. 2015) (Fig. S1).

### Transient expression of OsERS1–GFP in protoplasts and confocal fluorescence microscopy

Suspension cultures of rice Oc cell were maintained in 20 mL R2 S liquid medium (4.0 g/L KNO_3_, 335 mg/L (NH_4_)_2_ SO_4_, 273 mg/L NaH_2_ PO_4_, 166 mg/ L CaCl_2_ ·2H_2_O, 250 mg/L MgSO_4_ ·7H_2_O, 7.5mg/L Na_2_ EDTA, 5.5mg/L FeSO_4_ ·7H_2_O, 1.6 mg/L MnSO_4_ ·4H_2_O, 2.2 mg/L ZnSO_4_ ·7H_2_O, 3 mg/L H_3_ BO_4_, 0.125 mg/ L CuSO_4_ ·5H_2_O, 0.125 mg/L Na_2_ MoO_4_ ·2H_2_O, 30 g/L sucrose, 100 mg/L myo-inositol, 2 mg/L L-glycine, 1 mg/L thiamine-HCl, 0. 5 mg/L nicotinic acid, 0.5 mg/L pyridoxine-HCl, 2 mg/L 2,4-D, pH 5.8) at 110 rpm in the dark at 28°C. Cultures were subcultured weekly by transferring half of the cells to fresh media.

Protoplasts were prepared from cells 4 days post-transfer by incubating them in a digestion solution containing 1.5 % Cellulase RS (Yakult, Tokyo, Japan), 0.75% Macerozyme R-10 (Yakult), 0.6 M mannitol, 10 mM MES-KOH (pH 5.7), 10 mM CaCl_2_, and 1% BSA for 3 h in the dark with gentle shaking at 80 rpm. The protoplasts were then filtered through a 40 µm cell strainer and washed twice with W 5 solution (154 mM NaCl, 125 mM CaCl_2_, 5 mM KCl, and 2 mM MES-KOH at pH 7.5). Subsequently, they were suspended in W 5 solution at a concentration of 3–9 × 10^6^ cells/mL. To introduce GFP and RFP plasmids, 2.5 µg of each plasmid was added to 100 µL of the protoplast solution. A 110 µL PEG solution (40% PEG4000 (Fluka), 0.2 M mannitol, and 0.1M CaCl_2_) was then added, and the mixture was incubated at room temperature for 20 min. After incubation, 500 µL of W 5 solution was added, and the samples were gently inverted to mix. The protoplasts were pelleted by centrifugation at 800 rpm for 4 min, resuspended in 500 µL WI solution (0.2 M mannitol, 20 mM KCl, and 4 mM MES-KOH at pH 7.5), and incubated overnight in the dark at 28°C.

The GFP- and RFP-fusion proteins were observed using a confocal laser scanning microscope (D-ECLIPSE C1; Nikon). The GFP signal was detected using excitation at 488 nm with a 515/30 filter, while the RFP signal was detected using excitation at 543 nm with a 605/75 filter.

### Generation of the KO line by gene editing

To create KO lines of *OsERS1*, a 20-bp target sequence for single-guide RNA (sgRNA) was selected using CRISPR-P (http://cbi.hzau.edu.cn/crispr/). Oligonucleotides corresponding to the target sequence were annealed and cloned into the *Bbs* I site of the sgRNA cloning vector, pU6_ccdB_gRNA (Mikami, Toki and Endo 2015). The sgRNA expression cassette was excised with *Asc* I and *Pac* I and inserted into the CRISPR/Cas9 binary vector, pZH_gYSA_MMCas9 (Mikami, Toki and Endo 2015). The resulting T-DNA plasmid was used for the *Agrobacterium* -mediated transformation of rice (cv. Nipponbare) following the protocol described by (Toki 1997). The produced transgenic lines were grown on soil in a containment glasshouse at 28°C under a natural light condition to yield seeds. To identify *Osers1* mutant, genomic regions flanking the target sequence were amplified by PCR from T_0_ and T_1_ plants using the primers listed in Table S1. The amplified fragments were sequenced using Sanger sequencing. Homozygous mutants and nonmutant T_2_ plants were used for further study.

### Evaluation of the shoot ethylene response

Mature seeds were sown on filter paper in a Petri dish (9 cm diameter) containing 5 mL distilled water or 100 µM 1-aminocyclopropane-1 - carboxylic acid (ACC) solution. The seeds were incubated in the dark at 28°C for 7 days, after which the shoot (coleoptile and primary leaf) length was measured.

### Salt stress treatment

Mature seeds were sown on moist filter paper with 5 m L of distilled water as described above, and incubated in the dark at 28°C for 3 days. The seedlings were then transferred to new Petri dishes, with 10 seedlings per dish. Each dish received 6 mL of either distilled water or an NaCl solution (100 mM or 200 mM). The seedlings were incubated in the dark at 28°C for an additional 7 days. Shoot length was measured 10 days after imbibition (DAI).

### Expression analysis with quantitative RT-PCR (qRT-PCR)

Mature seeds were sown on moist filter paper with 5 mL distilled water and incubated in the dark at 28°C for 3 days. The seedlings were then transferred to new Petri dishes, with 10 seedlings per dish. Each dish received 8 mL of either distilled water or 100 µM ACC solution and was incubated in the dark at 28°C for 48 h. Shoots were collected at 5 DAI, immediately frozen in liquid nitrogen, and stored at −80°C. For each sample, three seedlings constituted a biological replicate, and four replicates were sampled.

Total RNA was prepared from the shoots using RNeasy Plant Mini Kit (Qiagen, Venlo, The Netherlands). Single-strand cDNA was synthesized from the total RNA using ReverTra Ace reverse transcriptase (Toyobo) and an oligo dT primer. Quantitative PCR was conducted with a real-time PCR cycler (Thermal Cycler Dice Real Time System Lite, Ta Ka Ra Bio Inc, Otsu, Japan) using the primers listed in Table S 1 and the KOD SYBR qPCR Mix (Toyobo). Four ethylene-responsive genes (*Germin-like*, Os08g0231400; *ERF063*, Os09g0287000; *ERF073*, Os09g0286600; kinase gene, Os07g0542600) were selected from a previous study (Ma 2013). The polyubiquitin gene served as a reference for normalizing transcript levels.

## Results

### Subcellular localization of OsERS1

Although ethylene receptors are primarily localized in the ER, a previous study reported that OsERS1 is localized in the plasma membrane when expressed in onion epidermal cells (Yu, Yau and Yip 2017). In this study, we examined the localization of OsERS1 by expressing a GFP-fusion protein in rice protoplasts. The entire CDS of *OsERS 1* was fused to the 5′-end of the GFP CDS to construct the *OsERS 1* – *GFP* fusion plasmid (Fig. S1). This construct, along with an RFP fusion construct of the ER marker TagRFP-AtSec22, was introduced into rice protoplasts. Confocal microscopy analysis revealed that OsERS1 –GFP signals were localized around the nucleus and dispersed in the cytoplasm (Fig. 1). Additionally, the GFP signals overlapped with the RFP signals from AtSec 22, indicating that OsERS1 is localized in the ER.

**Fig. 1.**
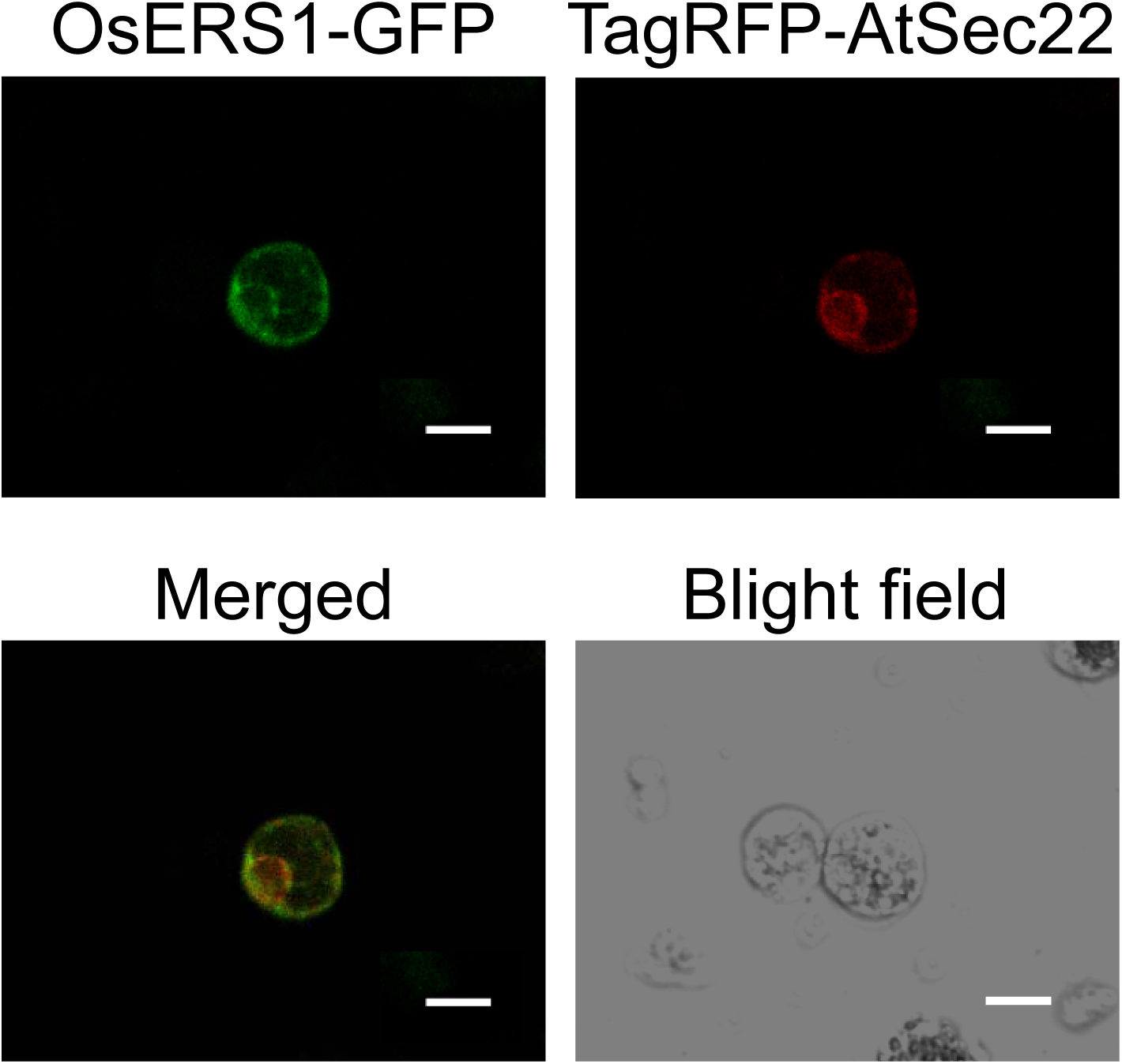
Subcellular localization of OsERS1. The OsERS1–GFP and Tag RFP-AtSec22 fusion proteins were transiently expressed in rice protoplasts. Scale bar = 10 μm.

### Generation of OsERS1 mutant lines by gene editing

To generate *OsERS1* KO mutants, an sgRNA targeting the 5′-end of the *OsERS1* CDS was designed (Fig. 2a). The sgRNA was cloned into a CRISPR/Cas9 gene-editing vector, and transgenic lines were produced by *Agrobacterium* -mediated transformation. Genomic sequencing of the target region in the resulting T_1_ plants revealed that three lines (N41, N61, and N152) contained a homozygous 1-bp insertion (Figs. 2b, 2c). This insertion caused a frameshift mutation, resulting in a premature stop codon that truncated the original 636-amino-acid protein after just 13 amino acids (Fig. 2d). This result suggested that the function of *OsERS1* was knocked out in these mutant lines. Conversely, two other transgenic lines (N21 and N131) showed no mutation in the target region and retained the wild-type (WT) sequence (Fig. 2c). These nonmutant lines served as controls in subsequent analyses.

**Fig. 2.**
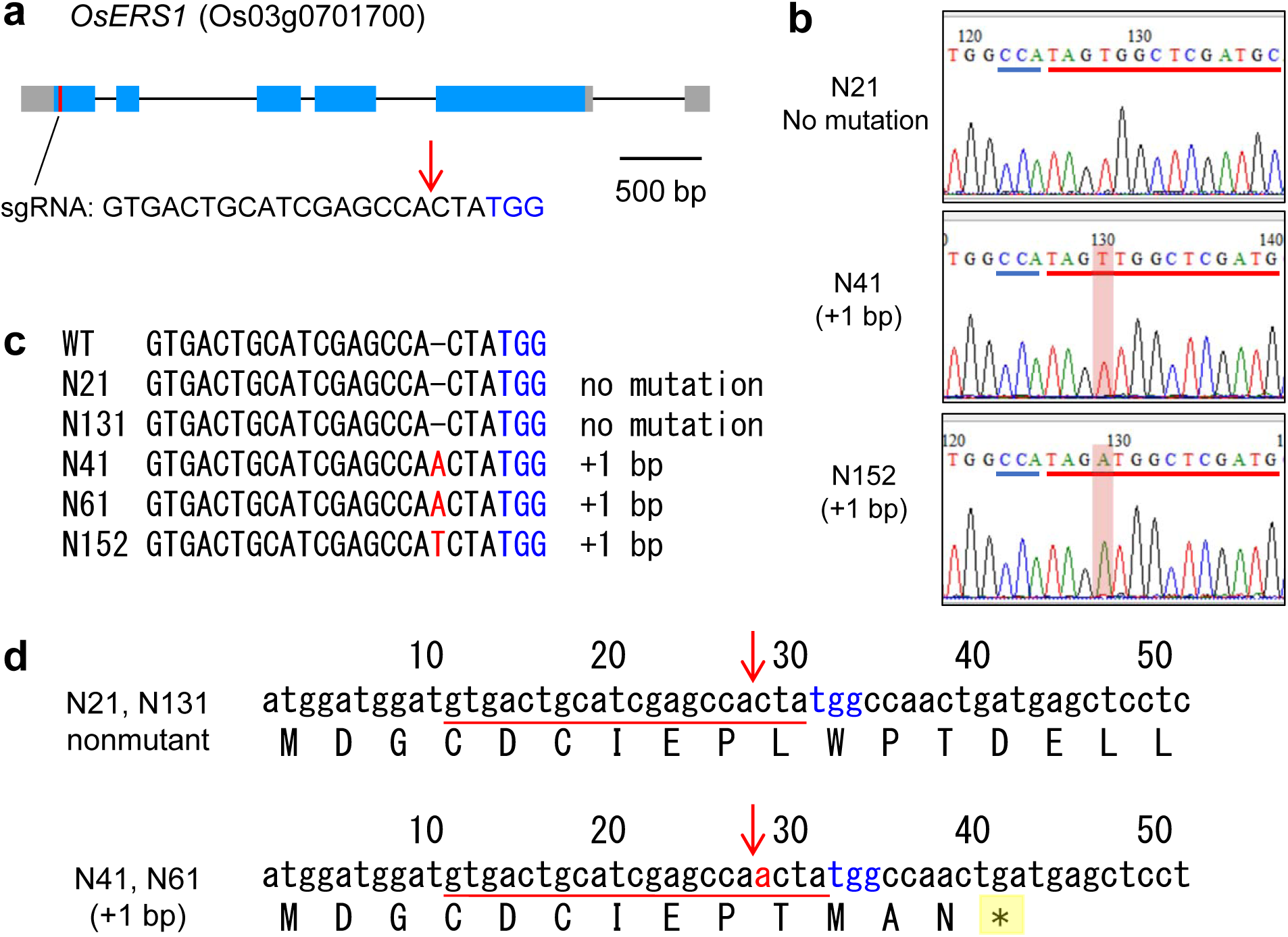
Generation of *OsERS1* -edited plants. (a) Design of sgRNA. (b–c) Genomic sequence of *OsERS1* in WT and mutant lines. (d) Nucleotide and deduced amino acid sequences in nonmutant and mutant lines. The blue letters in panels (a), (c), and (d) indicate the PAM sequence. The red lines in panels (b) and (d) highlight the target sequence. The vertical arrows in panels (a) and (d) indicate the cleavage site by Cas9.

### Enhanced shoot elongation in OsERS1 mutants

Ethylene promotes coleoptile elongation in etiolated seedlings, so we investigated whether ethylene-induced shoot elongation was affected in the *OsERS1* mutant lines (Fig. 3). We compared the shoot lengths of two nonmutant and three mutant lines and found that the mutant lines had significantly longer shoots than the nonmutant lines in the absence of ethylene. Treatment with an ethylene precursor, ACC, enhanced shoot elongation in both mutant and control lines. However, the mutant lines exhibited significantly longer shoots than the nonmutants. These results indicate that the *Os ERS1* mutation leads to enhanced shoot elongation both with and without ethylene.

**Fig. 3.**
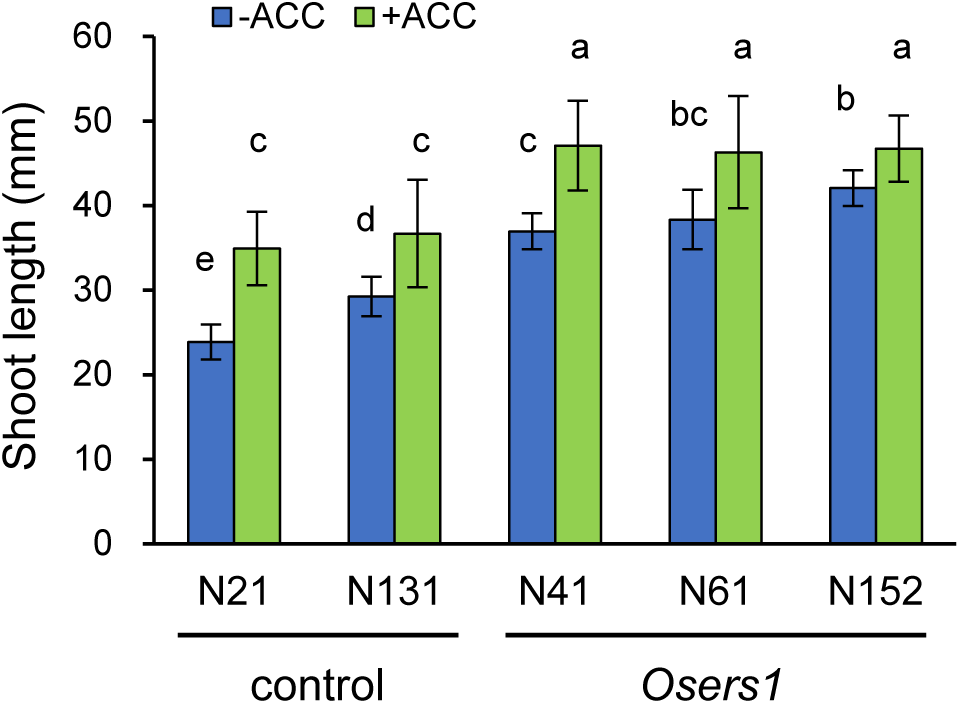
Enhanced shoot elongation in *OsERS1* KO lines. Etiolated seedlings of mutant and nonmutant lines were grown in the presence of 100 μM ACC for 7 days, and shoot length was measured. Data are presented as means ± SD (n = 24–30). Different letters indicate significant differences between samples (*P* < 0.05; Tukey–Kramer test).

### Enhanced expression of ethylene-responsive genes in OsERS1 mutants

As ethylene receptors negatively regulate the ethylene response, we hypothesized that the enhanced shoot elongation in the *OsERS1* mutants was caused by an elevated ethylene response due to *OsERS1* KO. To test this possibility, we analyzed the expression of ethylene-responsive genes as indicators of the ethylene response. The expression of four ethylene-responsive genes (*Germin-like*, *ERF063*, *ERF073*, and kinase gene) was measured in the N41 and N61 mutant lines, as well as the nonmutant line N131, using qRT-PCR. Two genes (*Germin-like* and kinase gene) exhibited similar or inconsistent expression patterns between the mutants and nonmutants (Fig. S2). However, the expression of two genes encoding ethylene-responsive transcription factors, *ERF063* and *ERF073*, was significantly higher in the mutant lines than in the control under ACC treatment, although expression levels were similar in the absence of ACC (Fig. 4). These findings suggest that the ethylene response is elevated in the *OsERS1* mutants when treated with ethylene.

**Fig. 4.**
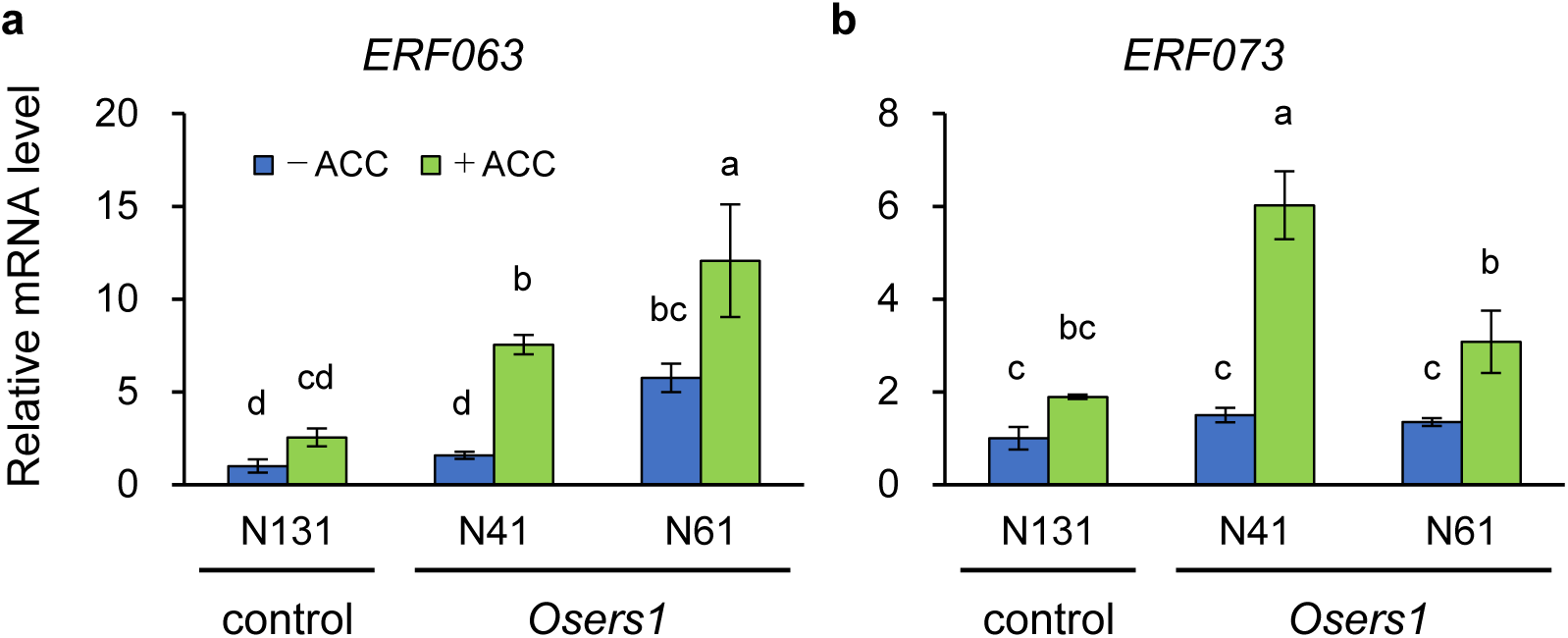
Expression of ethylene-responsive genes in *OsERS1* KO lines. Etiolated seedlings of mutant and nonmutant lines at 3 DAI were treated with 100 μM ACC for 48 h. Total RNA was prepared from the shoots of the seedlings, and qRT-PCR was conducted. Data are presented as means ± SD (n = 3–4). Different letters indicate significant differences between samples (*P* < 0.05; Tukey–Kramer test).

### Elevated salt tolerance in OsERS1 mutants

A previous study demonstrated that ethylene receptor isoforms have distinct roles in germination under salt stress in *Arabidopsis*. Specifically, ETR1 and EIN4 negatively regulate germination, while ETR2 has a positive effect (Wilson 2014). Therefore, we investigated whether the mutation of *OsERS 1* affects germination and seedling growth under salt stress. A germination test under 100–200 mM NaCl treatment for 2 days showed no significant difference in germination between the *OsERS1* mutant and nonmutant lines (data not shown). However, when we treated etiolated seedlings of the mutants with salt stress at post-germination stage, we found that the shoot length was significantly longer in the mutants than in the nonmutant lines under the treatment with 100 mM NaCl (Fig. 5). This result indicates that the loss of function of *OsERS1* elevated salt stress tolerance in rice seedlings.

**Fig. 5.**
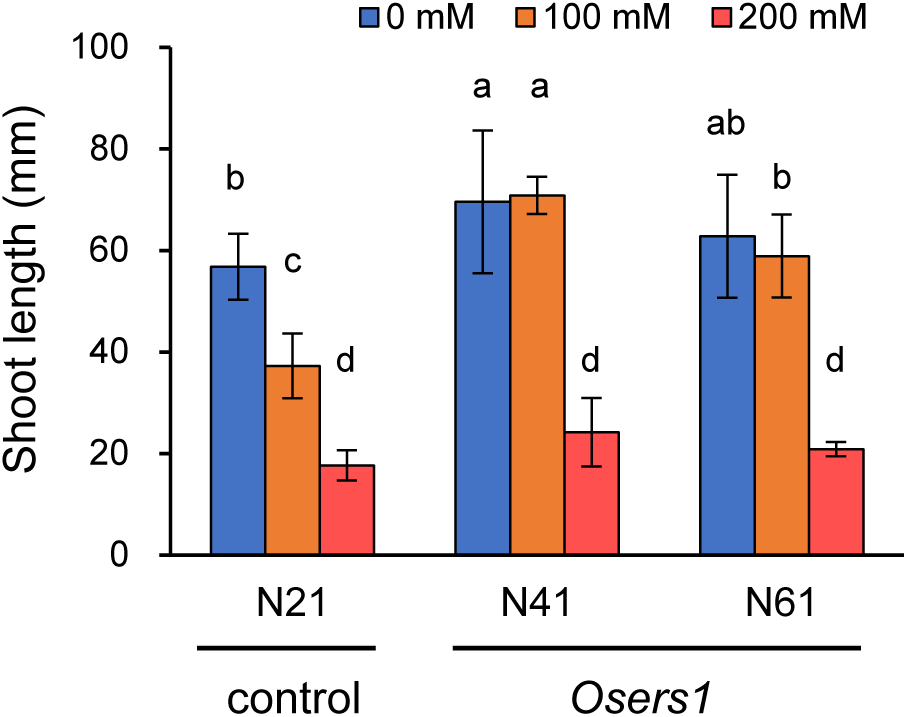
Enhanced salt tolerance in *OsERS1* KO lines. Etiolated seedlings of mutant and nonmutant lines at 3 DAI were treated with 100 or 200 mM NaCl for 7 days, and shoot length was measured. Data are presented as means ± SD (n = 10–19). Different letters indicate significant differences between samples (*P* < 0.05; Tukey–Kramer test).

## Discussion

In this study, we expressed an OsERS1–GFP fusion protein in rice protoplasts and observed its localization in the ER. This finding contrasts with a previous study in which OsERS1 was reported to localize in the plasma membrane (Yu, Yau and Yip 2017), where an N-terminal partial sequence of OsERS1 was fused to GFP and expressed in a heterologous system. Conversely, we analyzed the localization of a full-length OsERS1 by expressing it in rice protoplasts. Thus, our results indicate that OsERS1 is localized in the ER and likely functions there, similar to other ethylene receptors.

We also found that the KO of *OsERS1* resulted in enhanced shoot elongation in etiolated seedlings, both in the presence and absence of ethylene. This result suggests that *OsERS1* plays a role in regulating shoot growth. Additionally, the expression analysis of ethylene-responsive genes showed upregulation of ethylene-responsive transcription factors such as *ERF063* and *ERF073* in the *OsERS1* mutants. These findings suggest that the loss of function of *OsERS1* leads to an elevated ethylene response. Collectively, our results indicate that OsERS1 negatively regulates shoot elongation in etiolated seedlings. A previous study found that root growth in *OsERS1* mutant was suppressed in etiolated seedlings compared to WT plants, indicating elevated root ethylene response (Ma 2014). In conjunction with our findings, this suggests that OsERS1 suppresses the ethylene response in both shoots and roots. The loss of *OsERS 1* function leads to an ethylene response, resulting in enhanced shoot elongation and inhibited root growth.

Rice has five ethylene receptors, and this study suggests that Os ERS1 participates in regulating shoot elongation, as the KO of a single receptor resulted in a noticeable phenotype. However, KO mutants of *OsETR2*, *OsETR3*, and *OsERS2* also showed enhanced coleoptile growth in response to ethylene compared to WT plants (Wuriyanghan 2009), indicating that each receptor, including OsERS1, has a redundant role in regulating shoot (coleoptile) elongation. We observed that the *OsERS1* mutants exhibited enhanced shoot growth by ACC treatment, which is likely due to an amplified ethylene response mediated by other receptors in the absence of OsERS1.

The double mutant of *OsERS1* and *OsERS2* exhibits severe phenotypes, such as dwarfism and reduced seed setting under field conditions (Zhou 2020), suggesting that these receptors mainly function under light-grown conditions. However, our results, along with those from a previous study (Wuriyanghan 2009), indicate that four receptors, OsERS1, OsERS 2, OsETR2, and OsETR3 are involved in regulating growth during the early stages of etiolated seedling development. In our study, the expression patterns of two out of four ethylene-responsive genes remained unchanged between the *OsERS1* mutants and nonmutants, suggesting that each receptor regulates a distinct set of ethylene-responsive genes. Thus, these receptors play a redundant yet additive role in regulating shoot growth.

Ethylene is involved not only in growth regulation but also in stress response. In *Arabidopsis*, loss-of-function mutants of *EIN2*, *EIN3*, and *EIL1* show increased sensitivity to salt stress (Lei *et al*. 2011; Wang *et al*. 2007), indicating that ethylene signaling positively regulates salt tolerance (Riyazuddin 2020; Zhang 2016). Our result revealed that the *OsERS1* mutant exhibited improved shoot growth and enhanced tolerance under salt stress, suggesting that OsERS1 plays a role in salt stress tolerance. Given that ethylene signaling is likely activated in the *OsERS1* mutant, our findings imply that ethylene signaling positively regulates salt tolerance in rice seedlings. However, the exogenous ethylene treatment and overexpression of *OsEIL1* and *OsEIL2* have been shown to reduce salt tolerance in rice seedlings (Yang *et al*. 2015), suggesting that ethylene signaling negatively regulates salt stress response in rice, contrary to its role in *Arabidopsis*. Additionally, increased ethylene production caused by the overexpression of *OsARD1* enhances the expression of ABA biosynthesis genes and improves salt and drought stress tolerance (Liang *et al*. 2019). Thus, previous studies have shown contrasting results regarding the regulation of salt tolerance by ethylene. While green seedlings were used in both previous studies, we employed etiolated seedlings for salt stress treatment in this study. Thus, our result suggests that OsERS1 negatively regulates salt tolerance at least during the early growth stage in the dark.

Salt stress induces the production of reactive oxygen species (ROS) in plant cells, and in *Arabidopsis*, EIN3 and EIL1 regulate salt tolerance by suppressing ROS accumulation (Peng *et al*. 2014). OsEIL1 and OsEIL2 also regulate ROS scavenging genes in rice (Qiao *et al*. 2024). Exogenous ACC treatment enhances thermotolerance of rice seedlings by enhancing ROS scavenging capacity and upregulating heat shock factors and ethylene signaling components (Wu and Yang 2019). Therefore, the enhanced salt tolerance observed in the *OsERS 1* mutant may involve the upregulation of the ROS scavenging system through activated ethylene signaling.

In summary, this study found that OsERS1 is localized in the ER and participates in suppressing shoot growth in etiolated seedlings. During the post-germination growth stage, ethylene receptors, including Os ERS1, perceive ethylene, leading to enhanced shoot growth. Our data also indicate that the KO of *OsERS1*resulted in elevated salt tolerance in etiolated seedlings. The underlying mechanisms by which OsERS1 affects salt tolerance remain to be explored in future studies.

## Acknowledgments

The authors wish to thank Drs. Masaharu Kuroda and Masaki Endo at Institute of Agrobiological Sciences, NARO, Japan, for providing the ubiquitin promoter vector and the CRISPR/ Cas9 vector set, respectively. They also thank Dr. Tsutomu Kawasaki at Kindai University for providing the rice Oc cells.

## Supplementary material

Supplementary material is available at Bioscience, Biotechnology, and Biochemistry online.

## Data availability

All data generated for this study are included in the article.

## Author contribution

S.M. conceived the research and designed the experiments. S.M., S.T., Y. T., J.N., and M.H.S. conducted the experiments and analyzed the data. S.S. and T. M. supervised the study. S.M. wrote the manuscript.

