## Supplemental Data for "The rice ethylene receptor OsERS1 negatively regulates the shoot growth and salt tolerance in rice seedlings"

**Table S1.** Primers used in this study.

| Purpose and target (Gene ID) [1] | Primer name | Sequence [2] |
|---|---|---|
| Construction of the GFP-fusion plasmid | | |
| CDS of *OsERS1* | ERS1-G F1 | ATCTAGAGAGGACTGATGAACTTCTAAACATT |
| | ERS1-G R1 | ATACCATGGATCCAGATCCTATACTTCTCTGATACCGAGATTTC |
| Production of gene-editing lines of *OsERS1* | | |
| Construction of sgRNA | ERS1_N1_UP | gttgGTGACTGCATCGAGCCACTA |
| | ERS1_N1_DW | aaacTAGTGGCTCGATGCAGTCAC |
| Identification of mutation | ERS1_N1 Fw | TGTACTTCTGTAGGGAGTGTGGT |
| | ERS1_N1 Rv | GACTATAAATGCACCAAACTGGA |
| qRT-PCR for expression analysis | | |
| *Germin-like* (Os08g0231400) | Germin_F1 | GCTAATTGATTGGCTCCAATC |
| | Germin_R1 | TAGCAACATATCGTGACACAC |
| *ERF063* (Os09g0287000) | ERF063_F2 | AAGGCAAGGACCAACTTCCC |
| | ERF063_R2 | AGCAGCACTCCAGCAGTATG |
| *ERF073* (Os09g0286600) | ERF073_F2 | AGGGCCGCCAGCTTAATTAG |
| | ERF073_R2 | AATCGAGAGTCGCGGTAACC |
| Kinase gene (Os07g0542600) | Kinase_F1 | TCGACCTACAATGGCGGATG |
| | Kinase_R1 | CAAGCCAACAACGGTGCTAG |
| *Ubiquitin* (Os02g0161900) | Ubiquitin-1535 Fw | TTGTGTCTGGTTTAATGGACCATCGAGT |
| | Ubiquitin-1639 Rv | GGTTCAAATCCAACTTATTCATAAAGCA |

[1] Gene IDs are according to RAP-DB (https://rapdb.dna.affrc.go.jp).

[2] Lowercase letters indicate the additional sequences for insertion into *Bbs* I site.

**a**

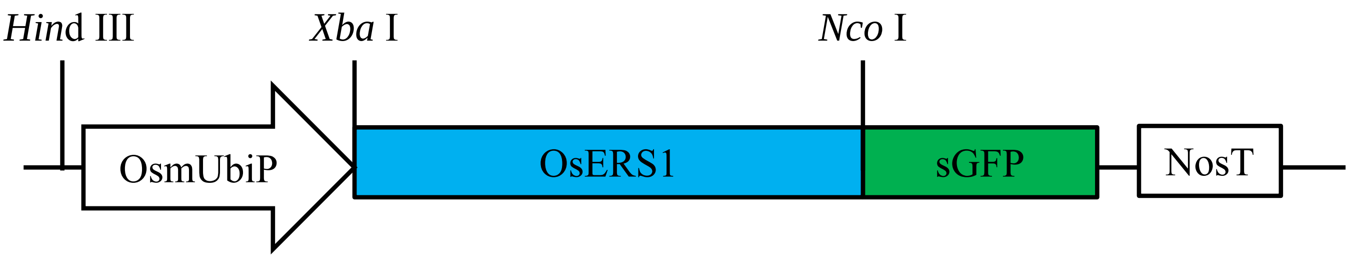


**b**

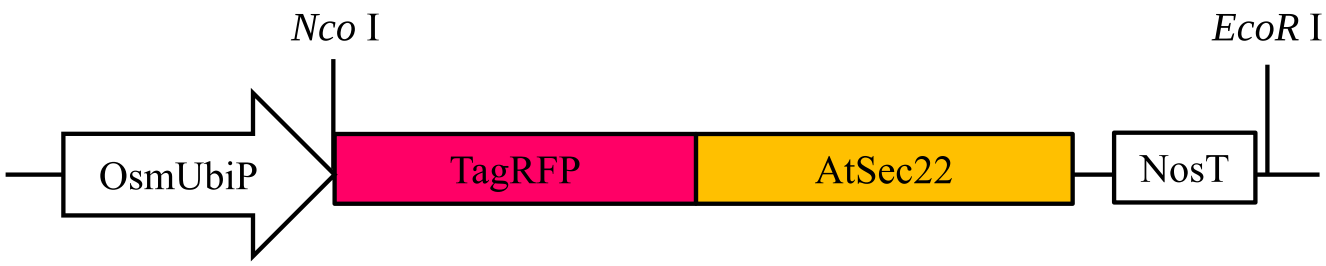


Supplementary Figure 1. GFP and RFP constructs for protein localization analysis. (a) OsERS1-GFP construct, (b) TagRFP-AtSec22 construct. OsmUbiP, the modified promoter of rice polyubiquitin gene; NosT, terminator of nopaline synthase gene.

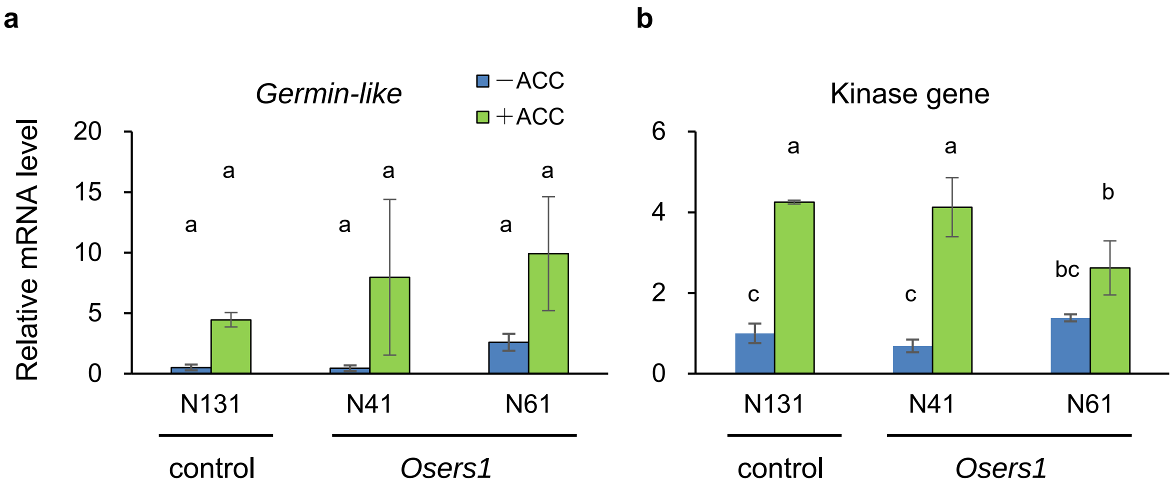


Supplementary Figure 2. Expression of ethylene-responsive genes in *OsERS1* KO lines. Etiolated seedlings of the mutant and nonmutant lines at 3 DAI were treated with 100 μM ACC for 48 h. Total RNA was prepared from the shoots of the seedlings, and qRT-PCR was conducted. Data are presented as means ± SD (n = 3–4). Different letters indicate significant differences between samples ($P < 0.05$; Tukey-Kramer tests).
